# Structural basis of nucleosome transcription mediated by Chd1 and FACT

**DOI:** 10.1101/2020.11.30.403857

**Authors:** Lucas Farnung, Moritz Ochmann, Maik Engeholm, Patrick Cramer

## Abstract

Transcription of eukaryotic protein-coding genes requires passage of RNA polymerase II (Pol II) through nucleosomes. Efficient Pol II passage through nucleosomes depends on the chromatin remodelling factor Chd1^1^ and the histone chaperone FACT^2^. How Chd1 and FACT mediate Pol II passage through nucleosomes remains unclear. Here we first show that Chd1 and FACT cooperate with the elongation factors Spt4/5 and TFIIS to facilitate Pol II transcription through a nucleosome in a defined biochemical system. We then determine cryo-EM structures of transcribing *Saccharomyces cerevisiae* Pol II-Spt4/5-nucleosome complexes with bound Chd1 or FACT at 2.9 Å and 3.1 Å resolution, respectively. In the first structure, transcribing Pol II has partially unwrapped nucleosomal DNA and exposed the proximal histone H2A/H2B dimer, which is bound by the acidic N-terminal region of Spt5 (Spt5N). The inhibitory DNA-binding region of Chd1 is released^3^ and the Chd1 translocase adopts an activated state that is poised to pump DNA towards Pol II. In the second structure, transcribing Pol II has generated a partially unravelled nucleosome that binds FACT in a manner that excludes Chd1 and Spt5N. These results suggest a dynamic model of Pol II passage through a nucleosome. In the model, Pol II enters the nucleosome^4^, activates Chd1 by releasing its DNA-binding region, and thereby stimulates its own progression. Pol II progression then enables FACT binding, liberates Chd1 and Spt5N, and eventually displaces a complex of FACT with histones that is transferred to upstream DNA.

---

Eukaryotic transcription occurs within chromatin. This poses the long-standing question of how Pol II passes through a nucleosome, the fundamental unit of chromatin^5,6^. Biochemical^4,7–9^ and single molecule assays^10,11^ have shown that Pol II pauses at distinct positions within the proximal half of the nucleosome^12–14^. Pol II enters the nucleosome and stably pauses just before it reaches the nucleosome dyad, at superhelical location (SHL) −1, both in vivo^15^ and in vitro^4,7,16^. To efficiently overcome the nucleosome barrier without loss of histones, Pol II requires the help of elongation factors^4,9,17–19^ ATP-dependent chromatin remodeling factors^20,21^ and histone chaperones^18,22,23^.

Recently, structures of Pol II-nucleosome complexes^4,9,16^ have provided a starting point to elucidate the mechanisms of chromatin transcription. These structures showed that transcription into the nucleosome results in partial DNA unwrapping. However, available structures lack the factors that are critical for nucleosome transcription, in particular the chromatin remodelling factor Chd1 and the histone chaperone FACT, a heterodimer of subunits Spt16 and Pob3. Chd1 and FACT form a complex^24,25^ and act in concert^26^. Binary structures of Chd1^27^ or FACT^28^ bound to nucleosomes revealed how these factors recognize their substrate. Here we use a structure-function analysis to investigate how Chd1 and FACT facilitate Pol II passage through a nucleosome and arrive at a mechanistic model of factor-mediated nucleosome transcription.

## Chd1 and FACT promote nucleosome transcription

To understand how Chd1 and FACT facilitate nucleosome passage by Pol II, we reconstituted factor-facilitated Pol II transcription through a nucleosome in vitro. We designed an extended nucleosome substrate for the biochemical and structural investigation of nucleosome transcription (Fig. 1a). The substrate consists of a single nucleosome, formed on a modified Widom 601 sequence^4^, with a 40-base pair (bp) upstream DNA extension (Methods). The DNA extension has a nine nucleotide (nt) 3’-overhang that enables annealing of a fluorescently labelled RNA oligonucleotide and allows for Pol II binding and catalytic RNA extension upon addition of nucleoside triphosphates (NTPs)^9^.

**Fig. 1:**
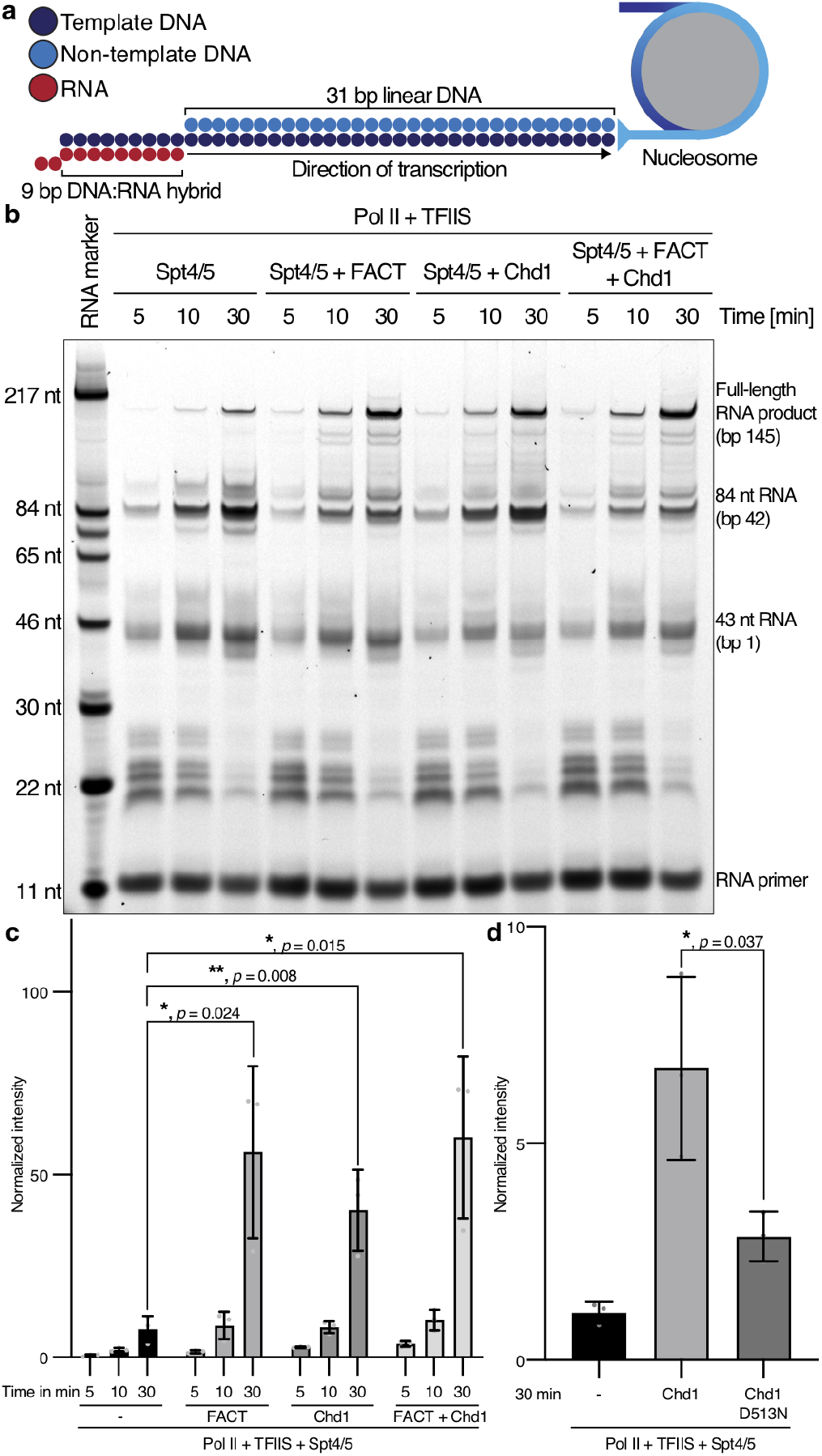
Chd1 and FACT stimulate nucleosome transcription. **a**, Schematic of nucleosome substrate used for formation of Pol Il-nucleosome complexes and RNA extension assays. **b**, Nucleosome transcription assay shows an increase in fulllength product in the presence of FACT, Chd1 or FACT and Chd1. RNA length and corresponding nucleosomal base pairs (bp) are indicated. **c**, Bar graph shows a significant increase in full-length product upon addition of FACT, Chd1 or FACT and Chd1 to Pol II-Spt4/5-TFIIS complexes after 30 min of transcription. n = 3 independent experiments with *P < 0.05, ** P < 0.01 with two-tailed t-test. **d**, Mutation of Chd1 to eliminate ATPase activity (D513N) strongly decreases production of full-length RNA product during nucleosomal transcription. n = 3 independent experiments with *P < 0.05, ** P < 0.01 with two-tailed t-test.

We then used a fluorescence-based RNA extension assay to determine the efficiency of nucleosome transcription in the presence of *S. cerevisiae* Pol II and various factors (Fig. 1b, Extended Data Fig. 1a). We formed a Pol II elongation complex on the extended nucleosome substrate, provided the elongation factors Spt4/5 and TFIIS, and initiated RNA elongation by the addition of 1 mM NTPs. Samples were removed at specific time points and the RNA products were separated by denaturing gel electrophoresis and quantified. We observed that Pol II paused upon nucleosome entry, at SHL −5 and at SHL −1 (Fig. 1b). A small fraction of Pol II could overcome these three barriers and transcribe trough the nucleosome, consistent with published observations^7,19^.

When Chd1 or FACT were individually added to the reactions containing Pol II, Spt4/5 and TFIIS, we observed a 5- or 7-fold increase in full-length product formation, respectively (Fig. 1b-c). The strong increase in full-length product in the presence of only Chd1 was dependent on the ATPase activity of Chd1, but some stimulatory effect was observed also with a catalytically inactive Chd1 variant (Fig. 1d, Extended Data Fig. 1b). Indeed, Chd1 binding to a nucleosome may facilitate transcription because it leads to detachment of two turns of DNA at the Pol II entry site^27^. The stimulatory effects of Chd1 or FACT depend on the presence of Spt4/5 (Extended Data Fig. 1c-d). When both Chd1 and FACT were included, full-length RNA product formation increased only slightly, suggesting that the stimulatory effects of Chd1 and FACT are not additive (Fig. 1b-c).

## Structure of Pol II-Spt4/5-nucleosome-Chd1 complex

To investigate the structural basis for Chd1 and FACT function during Pol II nucleosome passage, we formed a Pol II-Spt4/5-nucleosome complex in the presence of Chd1, FACT, and the transition state analogue ADP·BeF_3_ (Extended Data Fig. 2a-c). Transcription was carried out in the presence of GTP, CTP and UTP, and the complex was purified by size exclusion chromatography followed by mild crosslinking with glutaraldehyde (Methods). We prepared cryo-EM grids, collected a total of 3.76 million particles and obtained a reconstruction at a nominal resolution of 2.9 Å (Fig. 2, Extended Data Fig. 3, Table 1). We placed known structures and homology models of Pol II^29^, Spt4/5^30^ and the nucleosome core particle^31^ into the density, adjusted them, and modelled the remaining DNA (Extended Data Fig. 4). Additional density was observed for Chd1, but not for FACT, and was fitted with the structure of Chd1 in its post-translocated state^27^ (Extended Data Fig. 4h-I, 5a). The structure was real-space refined and has good stereochemistry (Table 1).

**Fig. 2:**
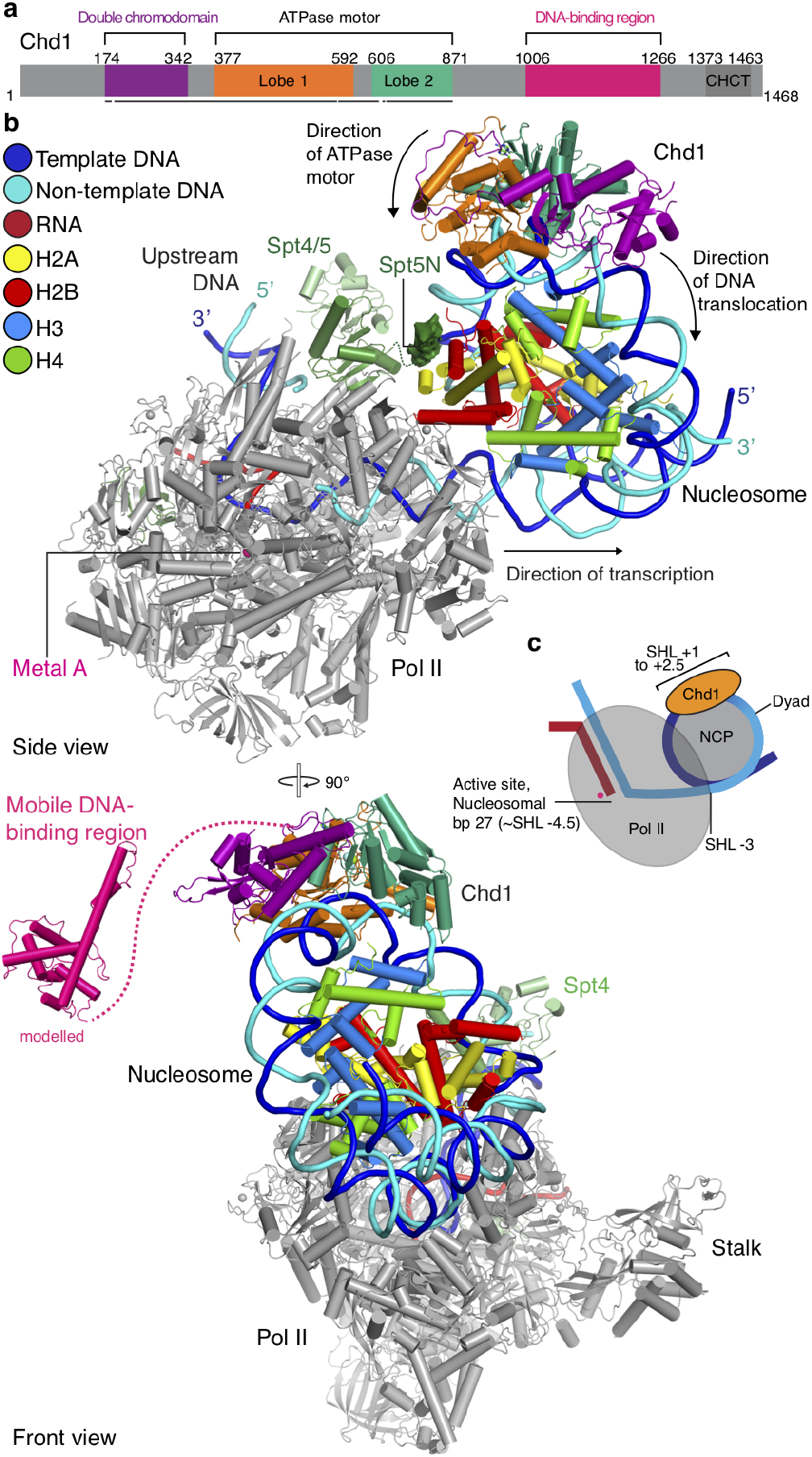
Pol II-Spt4/5-nucleosome-Chd1 structure. **a**, Chd1 domain architecture. Residues at domain boundaries are indicated. Regions modelled in the Pol II-Spt4/5-nucleosome-Chd1 structure are indicated with a black bar. Colour code used throughout. **b**, Two views of the structure related by a 90° rotation. Colour code for Pol II, Spt4/5, histones, metal A, RNA, template and non-template DNA is used throughout. Spt5N density is shown in surface representation. **c**, Schematic of the structure indicating key elements.

**Table 1:**
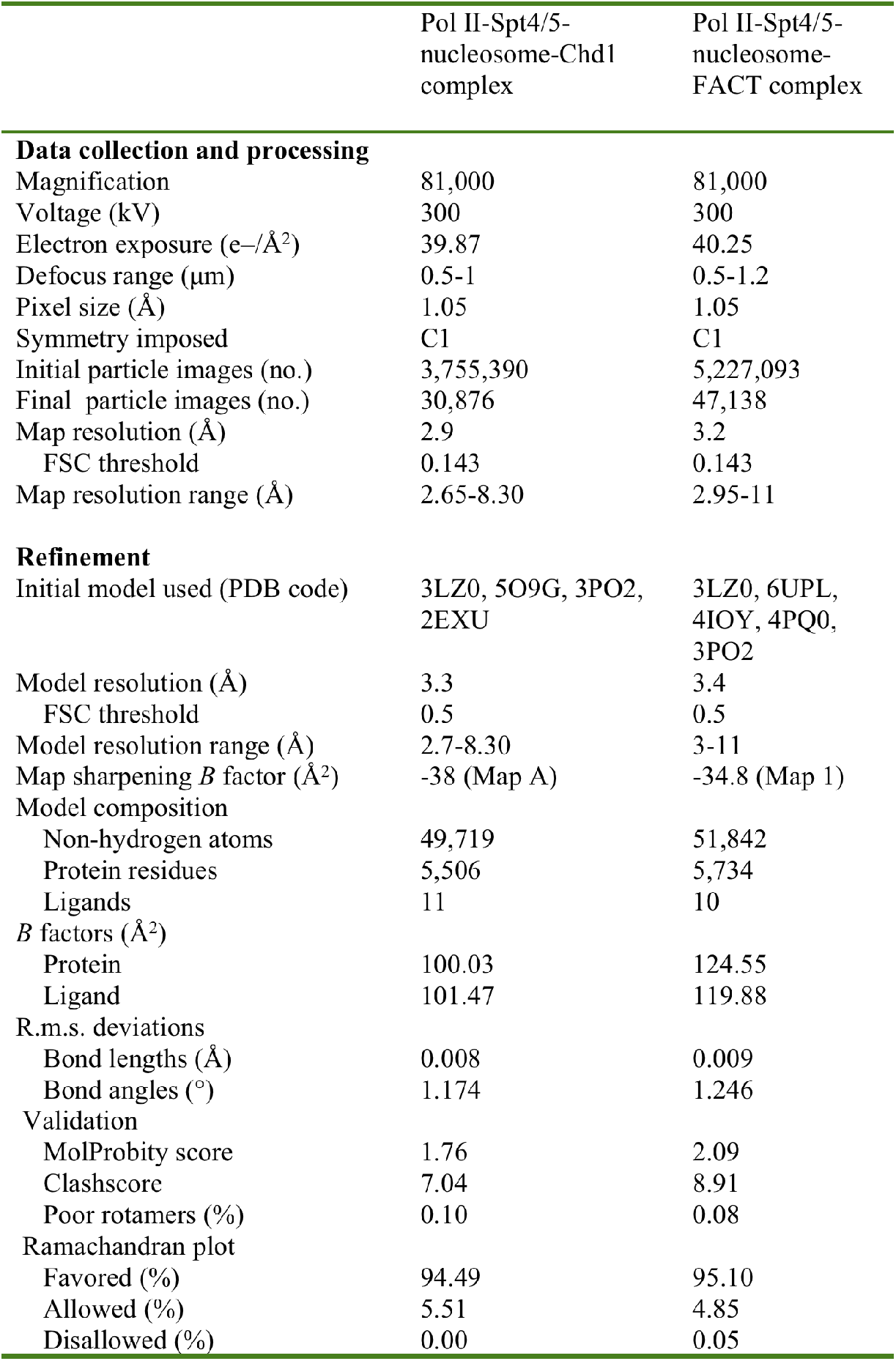
Cryo-EM data collection, refinement and validation statistics

|  | Pol II-Spt4/5-nucleosome-Chd1 complex | Pol II-Spt4/5-nucleosome-FACT complex |
| --- | --- | --- |
| **Data collection and processing** |  |  |
| Magnification | 81,000 | 81,000 |
| Voltage (kV) | 300 | 300 |
| Electron exposure (e<sup>-</sup>/Å<sup>2</sup>) | 39.87 | 40.25 |
| Defocus range (μm) | 0.5-1 | 0.5-1.2 |
| Pixel size (Å) | 1.05 | 1.05 |
| Symmetry imposed | C1 | C1 |
| Initial particle images (no.) | 3,755,390 | 5,227,093 |
| Final particle images (no.) | 30,876 | 47,138 |
| Map resolution (Å) | 2.9 | 3.2 |
| FSC threshold | 0.143 | 0.143 |
| Map resolution range (Å) | 2.65-8.30 | 2.95-11 |
| **Refinement** |  |  |
| Initial model used (PDB code) | 3LZ0, 5O9G, 3PO2, 2EXU | 3LZ0, 6UPL, 4IOY, 4PQ0, 3PO2 |
| Model resolution (Å) | 3.3 | 3.4 |
| FSC threshold | 0.5 | 0.5 |
| Model resolution range (Å) | 2.7-8.30 | 3-11 |
| Map sharpening *B* factor (Å<sup>2</sup>) | -38 (Map A) | -34.8 (Map 1) |
| Model composition |  |  |
| Non-hydrogen atoms | 49,719 | 51,842 |
| Protein residues | 5,506 | 5,734 |
| Ligands | 11 | 10 |
| *B* factors (Å<sup>2</sup>) |  |  |
| Protein | 100.03 | 124.55 |
| Ligand | 101.47 | 119.88 |
| R.m.s. deviations |  |  |
| Bond lengths (Å) | 0.008 | 0.009 |
| Bond angles (°) | 1.174 | 1.246 |
| Validation |  |  |
| MolProbity score | 1.76 | 2.09 |
| Clashscore | 7.04 | 8.91 |
| Poor rotamers (%) | 0.10 | 0.08 |
| Ramachandran plot |  |  |
| Favored (%) | 94.49 | 95.10 |
| Allowed (%) | 5.51 | 4.85 |
| Disallowed (%) | 0.00 | 0.05 |

The structure shows that Pol II adopts the active posttranslocated state and has transcribed 27 bp into the nucleosome, as observed biochemically (Extended Data Fig. 2c). Pol II has unwrapped 45 bp of nucleosomal DNA (Extended Data Fig. 4a, g). The Pol II front edge and active site are located around SHL –3 and –4.5, respectively (Fig. 2b-c, Extended Data Figs. 4a). Due to the unwrapping of DNA, the proximal histone H2A-H2B dimer is exposed and the nucleosome dyad faces away from Pol II. Spt4/5 binds around the Pol II cleft and the RNA exit channel as previously observed^32,33^. The NGN domain of Spt5 is sandwiched between upstream DNA and the downstream nucleosome. Some density is observed for the negatively charged N-terminal region of Spt5 (Spt5N). Spt5N is seen interacting with the exposed proximal H2A/H2B dimer, likely stabilizing it when DNA is absent (Fig. 2b, Extended Data Fig. 5b).

## Chd1 activation and Pol II progression

The structure shows that Chd1 binds the partially unwrapped nucleosome. Chd1 uses its double chromodomain and its ATPase motor domain to contact the nucleosome at SHLs +1 and +2, respectively (Fig. 2, Extended Data Fig. 4a). In contrast, the DNA-binding region of Chd1 is not observed in our structure and apparently mobile (Fig. 2b). The DNA-binding region is an inhibitory domain that restricts Chd1 ATPase activity when engaged with nucleosomal DNA^3^. We previously observed that the DNA-binding region binds the second DNA gyre in a nucleosome-Chd1 complex^27^ (Extended Data Fig. 5c). However, in the structure we present here, the second DNA gyre is no longer available for Chd1 interactions because DNA corresponding to SHL –7 to –5 has been transcribed by Pol II. These observations explain how Chd1 is activated during transcription elongation (Extended Data Fig. 5c). As Pol II transcribes into the nucleosome, it displaces the DNA-binding region of Chd1 from DNA. This releases the inhibitory effect of the DNA-binding region^3^ and activates Chd1, resulting in DNA translocation towards the nucleosome dyad and into the Pol II cleft^9^, thereby facilitating Pol II progression.

## Structure of Pol II-Spt4/5-nucleosome-FACT complex

To localize FACT during nucleosome transcription, we reconstituted a transcribing S. *cerevisiae* Pol II-Spt4/5-nucleosome-FACT complex by withholding Chd1 from the assembly (Extended Data Fig. 2d-f). We employed singleparticle cryo-EM to determine the structure of the complex at a nominal resolution of 3.1 Å (Methods), with densities for FACT at lower local resolutions (Extended Data Fig. 6). The high-resolution density observed around the Pol II active site allowed us to unambiguously define the nucleic acid sequence register and revealed that Pol II had stalled at bp +17 of the nucleosome (Extended Data Fig. 2f, Extended Data Fig. 7). As in the first structure, Pol II adopts the active posttranslocated state (Extended Data Fig. 7d).

In the structure, four turns of nucleosomal DNA (SHL −7 to SHL −3) are unwrapped from the histone octamer, exposing the proximal H2A/H2B dimer (Fig. 3). FACT embraces nucleosomal DNA near the dyad position at SHL +0.5 and contacts the template strand DNA backbone with the middle domain of subunit Spt16 at SHL −0.5 (Fig. 3b-c). Compared to the structure of the *H. sapiens* FACT-subnucleosome complex^28^, the position of FACT appears shifted towards the Pol II distal side of the nucleosome by one helical turn of DNA (Extended Data Fig. 8a). The previously observed position of FACT cannot be adopted in our structure because DNA is present on the distal side of the nucleosome, but is absent in the isolated FACT-subnucleosome complex structures that were reconstituted with shorter DNA^28^. In summary, our structure shows the position of FACT on a nucleosome that was partially unravelled by active Pol II transcription.

**Fig. 3:**
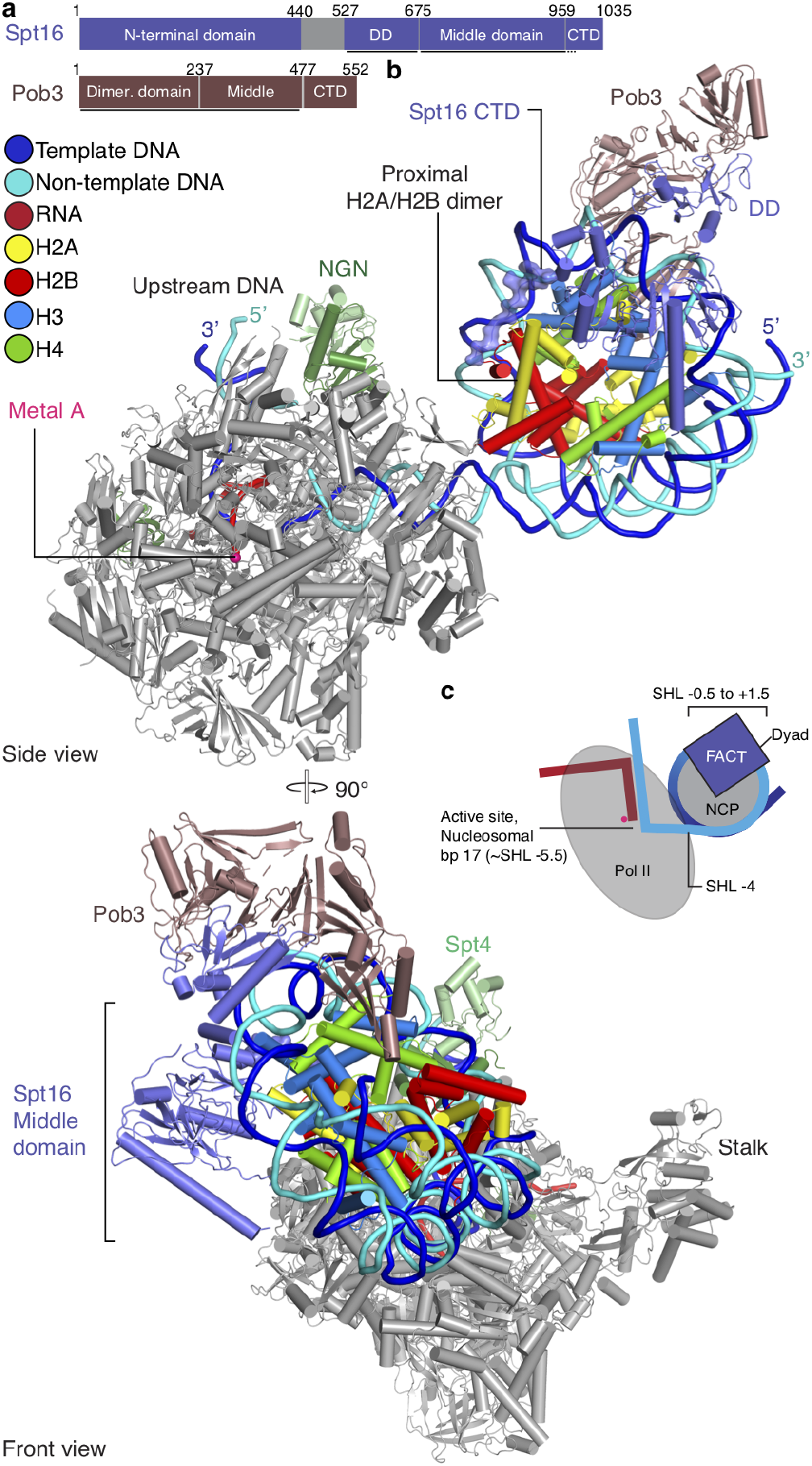
Pol II-Spt4/5-nucleosome-FACT structure. **a**, Domain architecture of FACT subunits Spt16 and Pob3 (DD, dimerization domain; CTD, C-terminal domain). Residues at domain boundaries are indicated. Regions modelled in the Pol II-Spt4/5-nucleosome-FACT structure are indicated with a black bar. **b**, Two views of the structure related by a 90° rotation. Colour code for Pol II, Spt4/Spt5, histones, Spt16, Pob3, metal A, RNA (red), template (blue) and non-template DNA (cyan) is used throughout. **c**, Schematic of the structure indicating key elements.

## Retention of the proximal H2A/H2B dimer

In our second structure, FACT additionally binds the exposed proximal H2A-H2B dimer using the C-terminal region of Spt16 (Fig. 3, Extended Data Fig. 7h-i), as previously observed^28,34^. This is consistent with the known role of FACT in the retention of the proximal H2A/H2B dimer during transcription^8,35,36^. Comparison with our first structure shows that the C-terminal region of Spt16 has replaced Spt5N for H2A/H2B binding. This suggests that Spt5N retains the exposed proximal H2A/H2B dimer until FACT is recruited. In our structure, FACT only binds the partially unwrapped nucleosome, and does not interact with Pol II or Spt4/5, consistent with recruitment of FACT to transcribed nucleosomes, rather than the transcription complex, in vivo^26,37^.

Superposition of our two structures results in a clash between Chd1 and the FACT subunit Pob3 (Extended Data Fig. 8b). This indicates that binding of Chd1 and FACT to the nucleosome is mutually exclusive, at least in the states we trapped here. Additionally, binding of the C-terminal domain of Spt16 to the H2A-H2B dimer is predicted to prevent reassociation of DNA with the exposed histone octamer surface, as had been observed in previous cryo-EM studies of Pol II-nucleosome complexes that revealed ‘foreign’ DNA of unknown origin^4^. These observations support a dynamic mechanism of subsequent or alternating Chd1 and FACT action.

## Model for nucleosome transcription

Our results converge with published data on a model for how Chd1 and FACT mediate nucleosome transcription (Extended Data Fig. 10a). When the Pol II-Spt4/5 complex transcribes into a Chd1-bound nucleosome, it releases the DNA-binding region of Chd1. This activates the Chd1 translocase^3^ and facilitates Pol II progression, which exposes the proximal H2A/H2B dimer that is temporarily stabilized by Spt5N. Pol II progression then generates a binding site for FACT, which displaces Chd1 and Spt5N. Modelling suggests a 30 bp window for FACT binding during Pol II progression (Extended Data Fig. 10b). Further Pol II progression would displace FACT from downstream DNA and enable FACT to transfer histones to upstream DNA or to relocate from the proximal to the distal H2A/H2B dimer, thereby preventing loss of histones^2,38^. Although the order of events during nucleosome transcription remains to be studied further, it appears that Chd1 functions upstream of FACT because Chd1 can bind complete nucleosomes, whereas FACT can only bind partially unravelled nucleosomes. Chd1 may even recruit FACT because it is known to interact with FACT^24,25^. Indeed, we could confirm the Chd1-FACT interaction biochemically and found that Chd1 binds FACT with its flexible N-terminal region (Extended Data Fig. 9). This is consistent with the idea that FACT is recruited near the nucleosome by Chd1 but remains flexible and only binds the nucleosome once DNA is partially unwrapped by transcription. In conclusion, we provide molecular snapshots of the dynamic process of factor-mediated nucleosome transcription and a model for Pol II progression through the nucleosome.

## ACKNOWLEDGEMENTS

We thank C. Oberthür and F Grabbe for help with protein purification, U. Neef and P. Rus for insect cell maintenance, and C. Dienemann and U. Steuerwald for maintenance of cryo-EM resources. We thank S.M. Vos for critical reading and input. M.E. is funded by the Deutsche Forschungsgemeinschaft (DFG, German Research Foundation) under Germany’s Excellence Strategy - EXC 2067/1-390729940. P.C. was supported by the Deutsche Forschungsgemeinschaft (SFB860, SPP2191, EXC 2067/1-390729940) and the ERC Advanced Investigator Grant CHROMATRANS (grant agreement No 693023).

## AUTHOR CONTRIBUTIONS

L. F. designed and conducted experiments and interpreted data, unless stated otherwise. M.O. assisted with purification of Spt4/5 and RNA extension assays. L.F. and M. O. prepared the Pol II-Spt4/5-nucleosome-Chd1 and Pol II-Spt4/5-nucleosome-FACT complexes for cryo-EM. M.E. conducted initial FACT-nucleosome binding experiments. P.C. supervised research. L.F. and P.C. wrote the manuscript with input from all authors.

## DATA AVAILABILITY STATEMENT

Reconstructions and structure coordinate files will be made available in the Electron Microscopy Database and the Protein Data Bank.

## Methods

No statistical methods were used to predetermine sample size. The experiments were not randomized, and the investigators were not blinded to allocation during experiments and outcome assessment.

### Molecular cloning

*Saccharomyces cerevisiae* Spt4 and Spt5 were cloned into vectors 438-A and 438-C, respectively, using ligation independent cloning (LIC). Vectors 438-A and 438-C were a gift from Scott Gradia, Addgene plasmids #55218 and #55220. Using LIC, the two genes were combined on a single 438-series vector. The construct contains Spt5 with an N-terminal 6× His tag followed by a maltose binding protein (MBP) tag, and a tobacco etch virus protease cleavage site. Spt4 does not contain tags. Each subunit in the combined vector is preceded by a PolH promoter and followed by a SV40 terminator. *Saccharomyces cerevisiae* Spt6 was cloned into vector 438-C using LIC. The construct contains an N-terminal 6× His tag followed by a maltose binding protein (MBP) tag and a tobacco etch virus protease cleavage site. A codon-optimized sequence of *S. cerevisiae* TFIIS for expression in *E. coli* was cloned into LIC-compatible vector 1-O. Vector 1-O was a gift from Scott Gradia, Addgene plasmid #29658. The construct contains an N-terminal 6× His tag followed by a Morc solubility tag and a tobacco etch virus protease cleavage site.

### Preparation of protein components

*S. cerevisiae* Pol II was purified as described^39^. *S. cerevisiae* Chd1 and FACT were expressed and purified as described^27^. All insect cell lines used for expression were purchased from Life Technologies (Sf9, Sf21) or from Expression Systems (Hi5) and used as identified by the vendor. Cell lines were not tested for mycoplasma contamination.

*S. cerevisiae* Spt4 and Spt5 were co-expressed in insect cells using a similar approach as reported previously^40^. After harvest, cell pellets were resuspended in lysis buffer 500 (500 mM NaCl, 20 mM Na·HEPES pH 7.4, 10% (v/v) glycerol, 1 mM DTT, 30 mM imidazole pH 8.0,0.284 μg ml^-1^ leupeptin, 1.37 μg m pepstatin A, 0.17 mg ml^-1^ PMSF, 0.33 mg ml^-1^ benzamidine). Cells were lysed by sonication. The cell lysate was applied to centrifugation (18,000g, 4 °C, 30 min) and ultracentrifugation (235,000g, 4 °C, 60 min). The supernatant containing Spt4/5 was subsequently filtered using 0.2 μm syringe filters (Millipore). The filtered supernatant was applied to a GE HisTrap 5 mL HP (GE Healthcare), pre-equilibrated in lysis buffer. The column was subsequently washed with 3 column volumes (CV) lysis buffer 500, 3 CV high salt buffer (1000 mM NaCl, 20 mM Na·HEPES pH 7.4, 10% (v/v) glycerol, 1 mM DTT, 30 mM imidazole pH 8.0, 0.284 μg ml^-1^ leupeptin, 1.37 μg ml^-1^ pepstatin A, 0.17 mg ml^-1^ PMSF, 0.33 mg ml^-1^ benzamidine) and 4.5 CV lysis buffer. Bound protein was eluted by gradient over 9 CV to 100 % nickel elution buffer (500 mM NaCl, 20 mM Na·HEPES pH 7.4, 10 % (v/v) glycerol, 1 mM DTT, 500 mM imidazole pH 8.0, 0.284 μg ml^-1^ leupeptin, 1.37 μg ml^-1^ pepstatin A, 0.17 mg ml^-1^ PMSF, 0.33 mg ml^-1^ benzamidine) over 9 CV. Fractions containing Spt4/5 were pooled and dialysed for 16 hours against dialysis buffer (300 mM NaCl, 20 mM Na·HEPES pH 7.4, 10 % (v/v) glycerol, 1 mM DTT, 30 mM imidazole pH 8.0, 0.284 μg ml^-1^ leupeptin, 1.37 μg ml^-1^ pepstatin A, 0.17 mg ml^-1^ PMSF, 0.33 mg ml^-1^ benzamidine). The dialyzed sample was applied to a GE HisTrap 5 mL HP and GE HTrap Q 5mL HP column combination. After application of the sample, the HisTrap 5 mL HP was removed and the HiTrap Q 5mL HP was washed with 5 CV dialysis buffer. Spt4/5 was eluted using a gradient elution to 100 % high salt buffer over 9 CV. Fractions containing Spt4/5 were concentrated using an Amicon Millipore 15 mL 50,000 MWCO centrifugal concentrator and applied to a GE Superdex 200 Increase 10/300 GL size exclusion column, pre-equilibrated in gel filtration buffer (500 mM NaCl, 20 mM Na·HEPES pH 7.4,10 % (v/v) glycerol, 1 mM DTT, 30 mM imidazole pH 8.0). Peak fractions were analysed with SDS-PAGE. Fractions containing Spt4/5 were concentrated using an Amicon Millipore 15 mL 50,000 MWCO centrifugal concentrator to a concentration of 20 μM, aliquoted, flash-frozen and stored at −80 °C. Typical preparations yield 300 μg of Spt4/5 from 1.2 L of insect cell culture. *S. cerevisiae* Spt6 was expressed in insect cells and subsequently purified with a similar protocol used for *H. sapiens* SPT6^41^ with a final concentration of 60 μM. Typical yields are 10 mg from 1.2 L of insect cell culture.

*S. cerevisiae* TFIIS was expressed in *E. coli* BL21 (DE3) RIL cells grown in LB medium. Cells were grown to an optical density at 600 nm of 0.6 at 37 °C. Expression of TFIIS was subsequently induced with 1 mM isopropyl β-D-1-thiogalactopyranoside at 18 °C for 16 hours. Cells were harvested by centrifugation and resuspended in TFIIS lysis buffer (500 mM NaCl, 20 mM Na·HEPES pH 7.4, 10 % (v/v) glycerol, 1 mM DTT, 30 mM imidazole pH 8.0, 0.284 μg ml^-1^ leupeptin, 1.37 μg ml^-1^ pepstatin A, 0.17 mg ml^-1^ PMSF, 0.33 mg ml^-1^ benzamidine). Cells were lysed by sonication. Two rounds of centrifugation (87,000g, 4 °C, 30 min) were used to clear the lysate. The supernatant was applied to a GE HisTrap 5 mL HP. Affinity purification, dialysis and TEV digest were performed similarly as described for Spt4/5. The dialyzed and TEV digested sample was applied to a GE HisTrap 5 mL HP, pre-equilibrated in TFIIS lysis buffer. The flow-through containing TFIIS was collected and concentrated using an Amicon Millipore 15 mL 10,000 MWCO centrifugal concentrator and applied to a GE S75 16/600 pg size exclusion column, pre-equilibrated in TFIIS size exclusion buffer (300 mM NaCl, 20 mM Na·HEPES pH 7.4,10 % (v/v) glycerol, 1 mM DTT, 30 mM imidazole pH 8.0). Fractions containing TFIIS were concentrated to 300 μM, aliquoted, flash frozen and stored at −80 °C. Typical yields are 10 mg from 6 L of *E. coli* expression culture. Protein identity of all protein components was confirmed by mass spectrometry.

### Nucleosome prepapration

*Xenopus laevis* histones were expressed and purified as described^27,42^. Histone H3 was H3K36Cme3 modified^43^. DNA fragments for nucleosome reconstitution were generated by PCR essentially as described27,36. A vector containing a modified Widom 601 sequence was used as a template for PCR. Super-helical locations are assigned based on previous publications^4,9,27,44^, assuming direction of transcription from negative to positive SHLs. Large-scale PCR reactions were performed with two PCR primers (forward primer: 5’-ACG AAG CGT AGC ATC ACT GTC TTG-3’’ reverse primer: 5’-ATC AGA ATC CCG GTG CCG AGG CCG C-3’) at a scale of 50 mL. The full-length PCR product (“Farnung construct”) has the following sequence: 5’-ACG AAG CGT AGC ATC ACT GTC TTG TGT TTG GTG TGT CTG GGT GGT GGC CGT TTT CGT TGT TTT TTT CTG TCT CGA ACC TGG AGA CTA GGG AGT AAT CCC CTT GGC GGT TAA AAC GCG GGG GAC AGC GCG TAC GTG CGT TTA AGC GGT GCT AGA GCT GTC TAC GAC CAA TTG AGC GGC CTC GGC ACC GGG ATT CTG AT-3’. PCR products were purified using anion exchange chromatography (GE Resoure Q 6mL) followed by EtOH precipitation. The DNA product was digested with TspRI in 1X CutSmart Buffer (NEB) overnight at 65 °C to generate the 9-nt ssDNA overhang. The digestion product was again purified with anion exchange chromatography, EtOH precipitated and resuspended in water. Nucleosome core particle reconstitution was then performed using the salt-gradient dialysis method^42^. The resulting nucleosome was purified using a Model 491 Prep Cell (Bio-Rad) and subsequently concentrated to 10-20 μM using an Amicon Millipore 15 mL 50,000 MWCO centrifugal concentrator. Quantification of the purified nucleosome was achieved by measuring absorbance at 280 nm. Molar extinction coefficients at 280 nm were determined for protein and nucleic acid components and were summed to yield a molar extinction coefficient for the reconstituted extended nucleosome.

### RNA extension assays

RNA extension assays were performed on the same nucleosomal template substrate used for the structural studies. A 6-FAM 5’ labelled 11 nt RNA (5’- /56-FAM/ rUrUrA rUrCrA rCrUrG rUrC-3’) was employed to monitor the transcription reaction. TFIIS has been added to all transcription reactions to prevent formation of overextended DNA·RNA hybrids and facilitate nucleosome passage^4,9^. All subsequent concentrations refer to the concentration in the final reaction. The final concentrations of buffer components are 130 mM NaCl, 20 mM Na·HEPES pH 7.4, 4 % (v/v) glycerol, 1 mM DTT/TCEP. The final volume for each RNA extension reaction is 40 μL. RNA (80 nM), nucleosomal template (80 nM) and *S. cerevisiae* Pol II (100 nM) were mixed in equimolar ratios and incubated for 5 minutes on ice. Spt4/5 (120 nM) and additional factors (500 nM each), 10× compensation buffer, and water were added to achieve final assay conditions. The sample was incubated for 3 minutes at 30 °C. Transcription elongation was started by the addition ATP, CTP, GTP and UTP (1 mM each) and TFIIS (60 nM). 5 μL of the reactions were quenched after 5 min, 10 min and 30 min in 5 μL 2× stop buffer (6.4 M urea, 50 mM EDTA pH 8.0, 1× TBE buffer) if time courses were performed. Samples were treated with 4 μg proteinase K for 15 minutes at 37 °C, denatured at 95 °C for 3 min, and separated by denaturing gel electrophoresis (4 μL of sample applied to an 8 M urea, 1× TBE buffer, 12 % acrylamide:bis-acrylamide 19:1 gel, run in 0.5× TBE buffer at 300 V for 30 minutes). RNA extension products were visualized using the 6-FAM label and a Typhoon 9500 FLA imager at an excitation wavelength of 473 nm and emission wavelength range of >520 nm.

Gels were subjected to linear contrast enhancement. All RNA extension assays were performed independently and at least three times. Full-length RNA extension products were quantified using Fiji 1.0. The products were normalized against the total intensity of the respective reaction lane to control for errors during gel loading. Bar charts show mean values and standard deviation as error bars. The following p values are applied * P < 0.05, ** P < 0.01. A two-tailed t-test was employed to determine statistical significance.

### Reconstitution of transcribing Pol II-nucleosome complexes

Complexes for cryo-EM were formed in a final buffer containing 130 mM NaCl, 20 mM Na·HEPES pH 7.4, 3 mM MgCl2, 1 mM DTT/TCEP, 4 % (v/v) glycerol. RNA (480 pmol, same construct as used for RNA extension assays) and nucleosome (120 pmol) were incubated for 5 min on ice. Pol II (120 pmol) Spt4/5 (180 pmol), Spt6 (180 pmol) was added and incubated for 5 min on ice. H2O and compensation buffer were added to reach final buffer conditions and the sample was incubated for 5 min. Transcription elongation was started by the addition of 1 mM of GTP, CTP, UTP each, and 0.4 mM 3’-dATP in the case of the Pol II-Spt4/5-nucleosome-FACT complex. Instead of 3’-dATP, 1 mM ADP·BeF_3_ was added to the Pol II-Spt4/5-nucleosome-Chd1-FACT complex. 108 pmol of TFIIS were immediately added after NTP addition.

After 15 minutes of incubation at 30 °C, Chd1 (180 pmol) and FACT (180 pmol), preincubated with H2A/H2B dimer (180 pmol), or FACT alone (180 pmol), preincubated with H2A/H2B dimer (180 pmol), were added. The transcription reactions were allowed to proceed for an additional 30 minutes at 30 °C and quenched with EDTA (10 mM final concentration). The samples were subsequently centrifuged and applied onto a Superose 6 3.2/300 Increase size exlusion column (GE Healthcare), equilibrated in complex buffer (100 mM NaCl, 20 mM Na·HEPES pH 7.4, 3 mM MgCl2, 1 mM TCEP, 4 % (v/v) glycerol). Fractions eluting from the size exclusion column were analysed using SDS-PAGE and denaturing urea PAGE. Consistent with previous observations (data not shown), TFIIS is lost from the elongation complex during size exlusion chromatography. Fractions containing the complex were individually cross-linked and dialyzed against dialysis buffer (100 mM NaCl, 20 mM Na·HEPES pH 7.4, 3 mM MgCl2, 1 mM TCEP) for three hours as described previously^45^.

The dialyzed complexes with an approximate concentration of 50-100 nM were applied to R2/2 gold grids, Au 200 mesh (Quantifoil). The grids were glow-discharged for 100 s prior to application of 2 μL of sample to each side of the grid. After incubation of the sample for eight seconds, the grid was blotted and vitrified by plunging into liquid ethane using a Vitrobot Mark IV (Thermo Fisher). The Vitrobot was operated at 4 °C and 100 % humidity. A blot force of 5 and blot time between 3-5 seconds were used for the sample preparation. The grids were clipped and subsequently stored in liquid nitrogen before data acquisition.

### Cryo-EM analysis and image processing

Cryo-EM data was collected on a Titan Krios II transmission electron microscope (FEI) operated at 300 keV. A K3 summit direct detector (GATAN) with a GIF Quantum Filter with a slit width of 20 eV was used for the data acquisition. Data acquisition was performed using SerialEM at a nominal magnification of 81,000×, corresponding to a pixel size of 1.05 Å/pixel in nanoprobe EF-TEM mode. For the Pol II-Spt4/5-nucleosome-Chd1 dataset, image stacks of 40 frames were collected in counting mode over 2.2 s. The dose rate was 18.12 e^-^ per Å^2^ per s for a total dose of 39.87 e^-^ per Å^2^. For the Pol II-Spt4/5-nucleosome-FACT dataset, image stacks of 40 frames were collected in counting mode over 2.2 s. The dose rate was 18.30 e^-^ per Å^2^ per s for a total dose of 40.25 e^-^ per Å^2^.

Micrographs were stacked and processed using Warp^46^. CTF estimation and motion correction was performed using Warp. Particles were picked using an in-house trained instance of the neural network BoxNet2 as implemented in Warp, yielding 3,755,390 particles for the Pol II-Spt4/5-nucleosome-Chd1 dataset and 5,227,093 particles for the Pol II-Spt4/5- nucleosome-FACT dataset. Particles were extracted with a box size of 400 pixels and normalized. Further image processing was performed with cryoSPARC^47^ and RELION 3.0.7^48^.

For the Pol II-Spt4/5-nucleosome-Chd1 dataset, particles were separated into two subsets and subsequently 3D classified using cryoSPARC. Particles were selected for the presence of a nucleosome-like density. Selected particles were imported into RELION. Two subsequent rounds of 3D classification resulted in 247,604 particles with clear nucleosomal density. A 3D refinement of these particles resulted in an overall model of 2.6 Å. The particles were CTF refined and Bayesian polishing was conducted. A mask encompassing the nucleosome and additional density at SHL +2 was applied to two rounds of 3D classification to select for particles that contain Chd1. This resulted in 30,876 particles with clear density for Chd1 bound to the partially unwrapped nucleosome. Particles were then subsequently 3D refined resulting in map A with a resolution of 2.9 Å (FSC threshold 0.143 criterion). The map showed excellent density for Pol II but the nucleosome-Chd1 part of the map showed flexibility. Therefore, Pol II with Spt4/5 and the nucleosome-Chd1 parts of the maps were individually refined using a mask for Pol II-Spt4/5 or nucleosome-Chd1, respectively. Signal subtraction was applied. This resulted in two masked refinements (Pol II-Spt4/5, map B; nucleosome-Chd1 with signal subtraction, map C) with overall improved density. These two masked refinements were combined using the Frankenmap and Noise2map tool set of Warp, resulting in the final composite map (map D).

For the Pol II-Spt4/5-nucleosome-FACT dataset, particles were separated into two subsets and subsequently 3D classified using cryoSPARC and RELION. Particles were selected for the presence of nucleosome-like densities. The selection resulted in 603,550 particles with clear nucleosome-like density. The selected particles were 3D refinement using an angular sampling rate of 7.5° and subjected to further classifications to select for particles with bound FACT. This resulted ultimately in 48,718 particles. These particles were again subjected to 3D classification. After 3D refinement, CTF re-finement and Bayesian polishing the remaining 47,138 particles resulted in a final refinement (map 1) with an overall resolution of 3.1 Å (FSC threshold 0.143 criterion). To improve densities for the nucleosome-FACT and Pol II-Spt4/5 parts of the map, the particles were subjected to masked refinements (maps 2 and 3). Similar to the Chd1 dataset, the masked refinements were combined using the Frankenmap and Noise2map tools included in Warp, yielding the final map (composite map 4). Local resolutions of the composite maps were estimated using the RELION built-in tool for the determination of local resolutions.

### Model building and refinement

For the Pol II-Spt4/5- nucleosome-Chd1 structure, structures of *S. cerevisiae* Pol II (PDB code 3PO2), X. laevis nucleosome (PDB code 3LZ0), Chd1 with ADP·BeF3 (PDB code 5O9G), and Spt4/5 (PDB codes 2EXU) were rigid body docked into map D and refined using COOT^49^. DNA from the elongation complex and nucleosomal DNA were connected using COOT. Density in the active site of Pol II allowed unambiguous assignment of DNA register. Surprisingly, the complex had transcribed over the T-less cassette that should stall further elongation. The ADP (Sigma-Aldrich) used in the formation of ADP·BeF3 is reported to be contaminated with up to 2.76 % ATP^50^, possibly providing the required substrate to transcribe past the end of the T-less cassette. Identification of DNA-RNA register was aided by map B.

For the Pol II-Spt4/5-nucleosome-FACT structure, the refined Pol II part of the Pol II-Spt4/5-nucleosome-Chd1 structure was rigid-body docked into the density. Additionally, *X. laevis* nucleosome (PDB code 3LZ0), *H. sapiens* FACT (PDB code 6UPK), and Spt4/5 (PDB codes 2EXU) were rigid body docked into map 4 using COOT. *S. cerevisiae* FACT structures (Spt16, PDB codes 4IOY; Pob3, 4PQ0) were then superposed onto the docked *H. sapiens* FACT structure. The dimerization domain of Spt16 and Pob3 were generated using PHYRE2^51^ and also superposed onto the docked FACT structure without any additional manual manipulation. The CTD of Spt16 was modelled de novo as a poly-alanine extension of Spt16 with no sequence assignment. DNA from the Pol II part of the structure and the nucleosomal DNA were connected in COOT. The density in the active site allowed for unambiguous assignment of the DNA register and the nucleic acid sequence adjusted accordingly. Identification of DNA-RNA register was aided by map 2.

Both atomic models were real-space refined using PHENIX^52^ with secondary structure restraints against map D (Pol II-Spt4/5-nucleosome-Chd1 model) and map 4 (Pol II-Spt4/5-nucleosome-FACT model).

### Figure generation

Figures for structural models were generated using PyMol (version 2.3.4) and UCSF ChimeraX^53^. Gel quantification was performed using Fiji and graphs were generated using Graphpad Prism.

---

**Extended Data Fig. 1:**
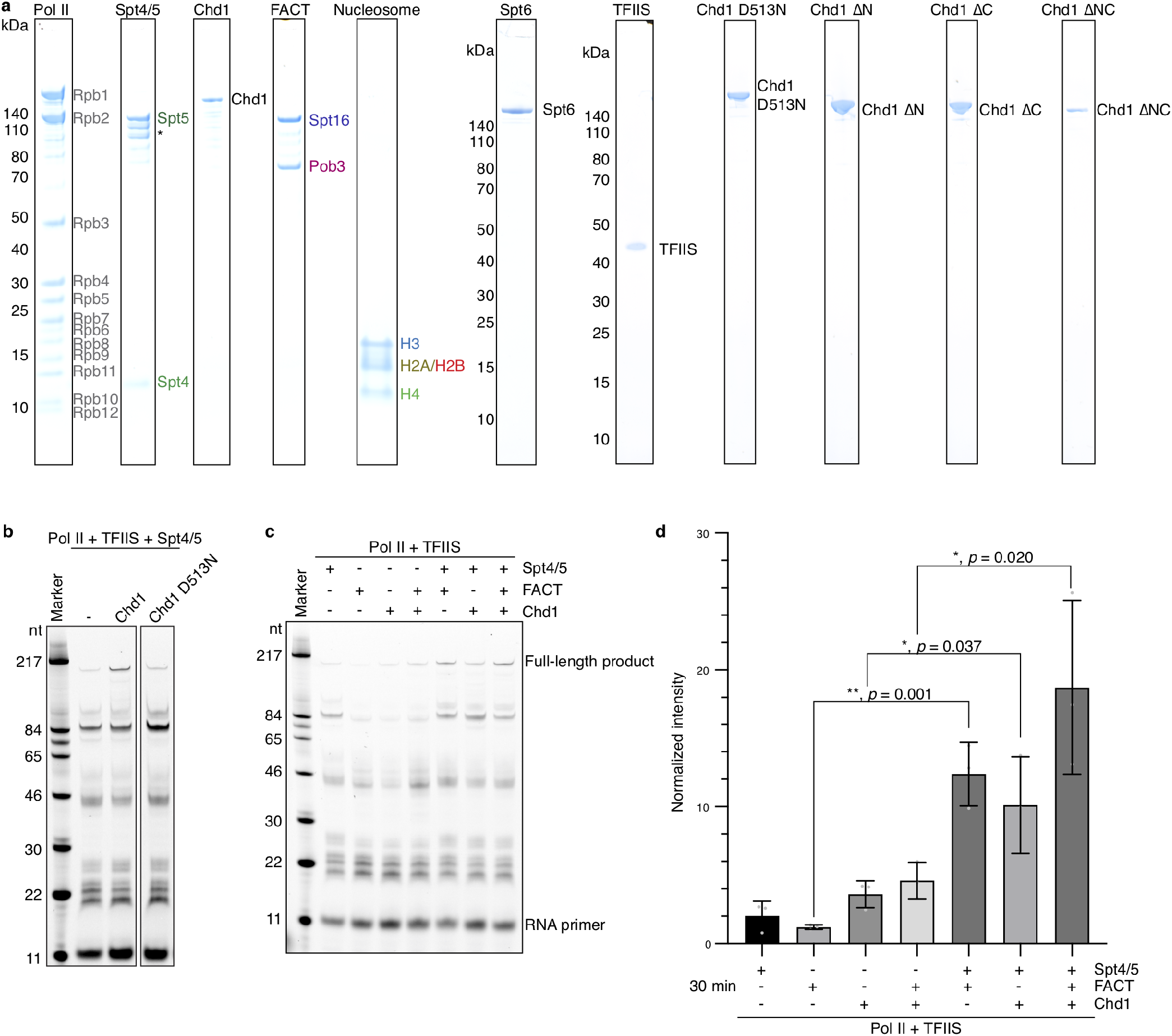
Additional information on RNA extension assays. **a**, SDS-PAGE of purified proteins. Purified proteins were run on 4-12 % Bis-Tris SDS-PAGE gels in 1X MES Buffer, stained with Coomassie Blue. Asterisk (*) demarcates degradation products of Spt5. **b**, 12 % denaturing urea gel of RNA extension assay with Chd1 D513N mutant. RNA is visualized using 6-FAM label on 5’ end of RNA. **c**, 12 % denaturing urea gel of RNA extension assay with different factor combinations reveals dependence of Spt4/5 for increase in full-length product in the presence of FACT and Chd1. RNA is visualized using 6-FAM label on 5’ end of RNA. **d**, Bar plot with quantification of c. Mean normalized intensity is shown as bar plot. Error bars represent standard deviation. Quantification from n = 3 independent experiments with *P < 0.05, ** P < 0.01 with two-tailed t-test.

**Extended Data Fig. 2:**
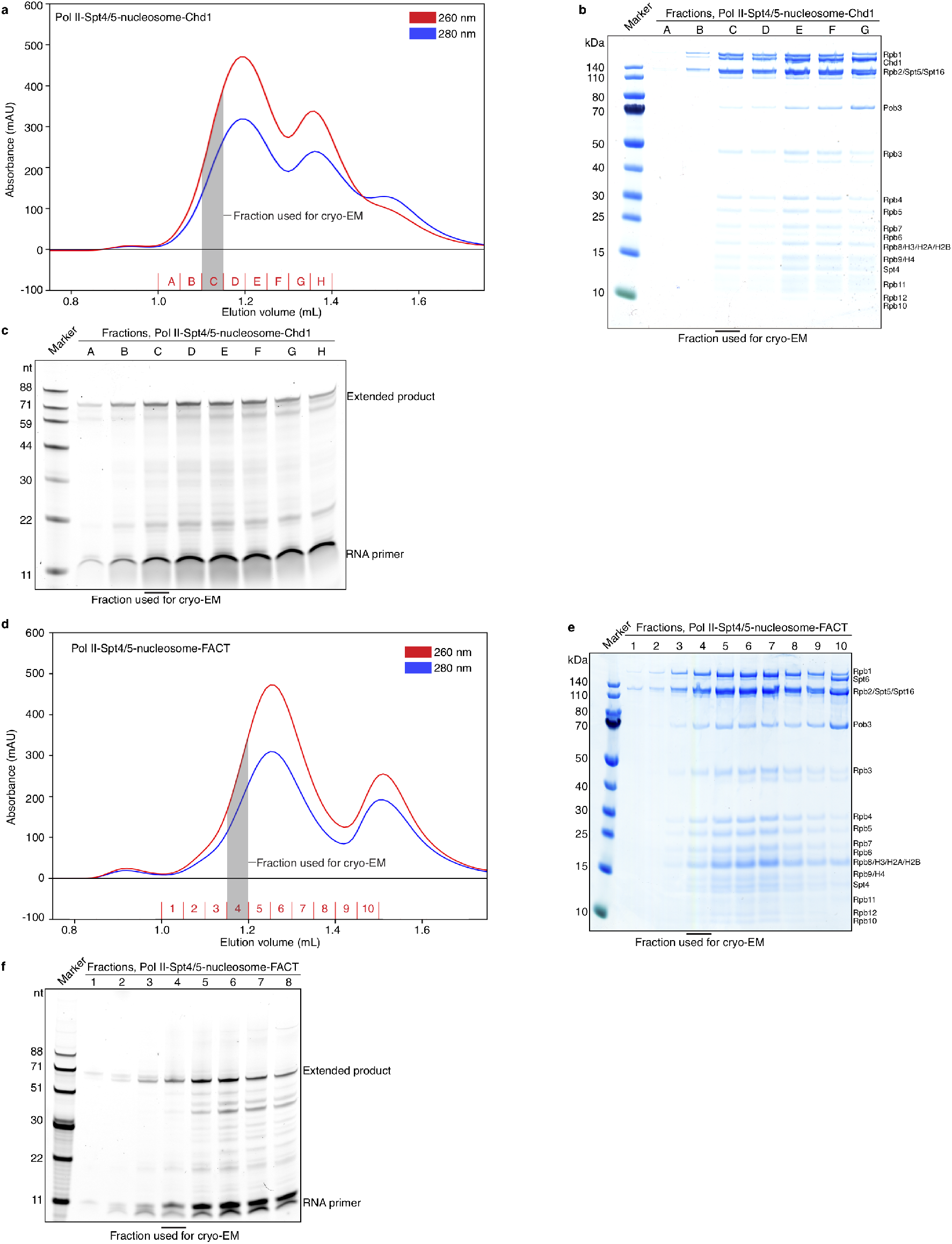
Formation of Pol II-nucleosome complexes with bound Chd1 and FACT. **a**, Chromatogram of Pol II-Spt4/5-nucleosome-Chd1 complex formation using size exclusion chromatography. Fractions used in further analysis are indicated. **b**, SDS-PAGE of Pol II-Spt4/5-nucleosome-Chd1 complex formation. SDS-PAGE shows presence of FACT in the complex. **c**, 12 % denaturing urea gel of Pol II-Spt4/5-nucleosome-Chd1 complex formation. RNA is visualized using 6-FAM label on 5’ end of RNA. **d**, Chromatogram of Pol II-Spt4/5-nucleosome-FACT complex formation using size exclusion chromatography. **e**, SDS-PAGE of Pol II-Spt4/5-nucleosome-FACT complex formation. **f**, 12 % denaturing urea gel of Pol II-Spt4/5-nucleosome-FACT complex formation. RNA is visualized using 6-FAM label on 5’ end of RNA.

**Extended Data Fig. 3:**
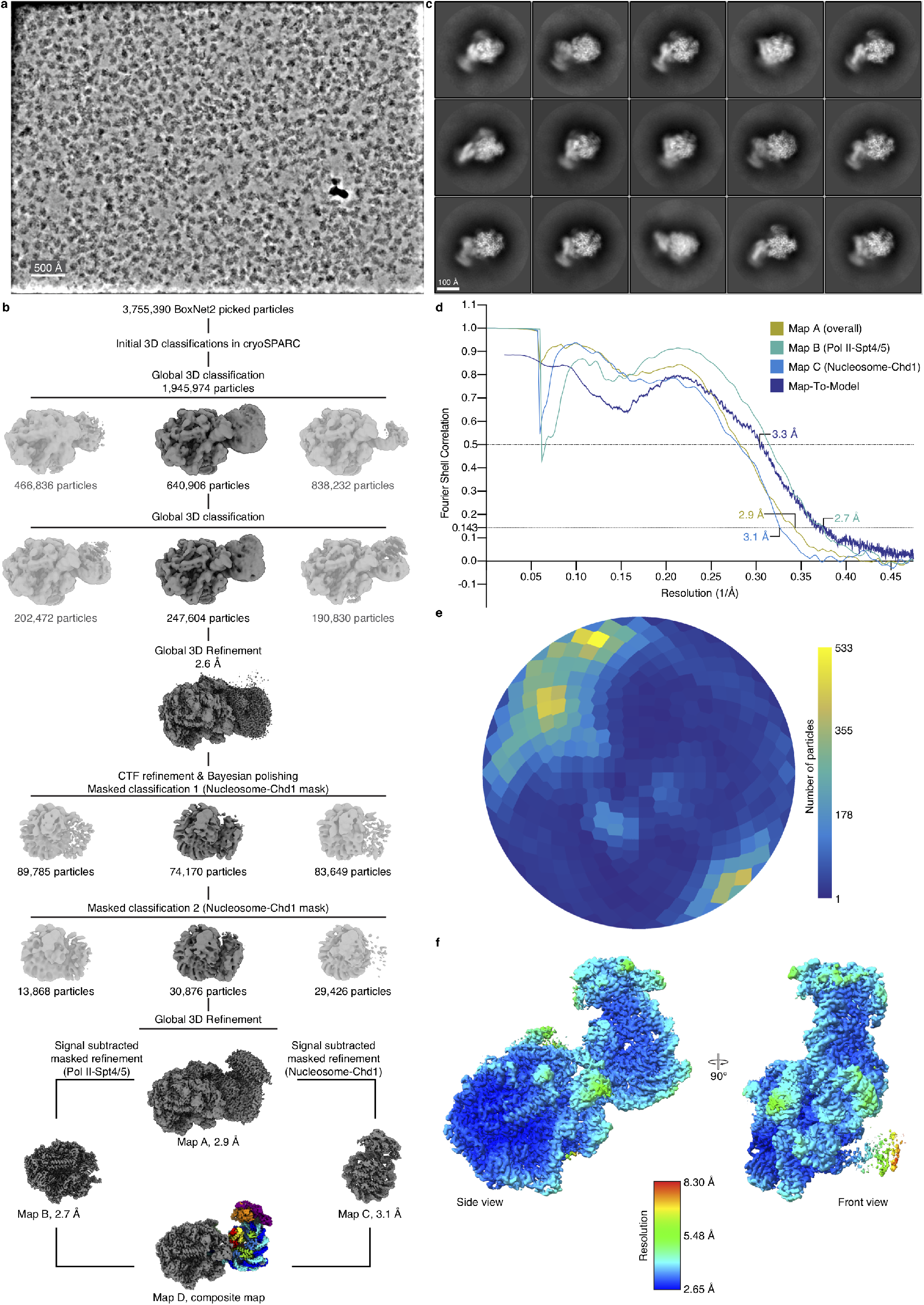
Data acquisition, processing, and data quality metrics for the Pol II-Spt4/5-nucleosome-Chd1 structure. **a**, Representative denoised micrograph of data collection with scale bar (50 nm). **b**, Sorting and classification tree of Pol II-Spt4/5-nucleosome-Chd1 dataset. **c**, 2D classes of final refinement show RNA polymerase II and nucleosome-like shape with additional density (Chd1) with scale bar of 10 nm. **d**, FSC curves of maps A-C and map-to-model. Resolutions at FSC threshold criterions 0.143 and 0.5 are indicated. **e**, Angular distribution of particles employed to reconstruct map A. **f**, Local resolution of composite map D.

**Extended Data Fig. 4:**
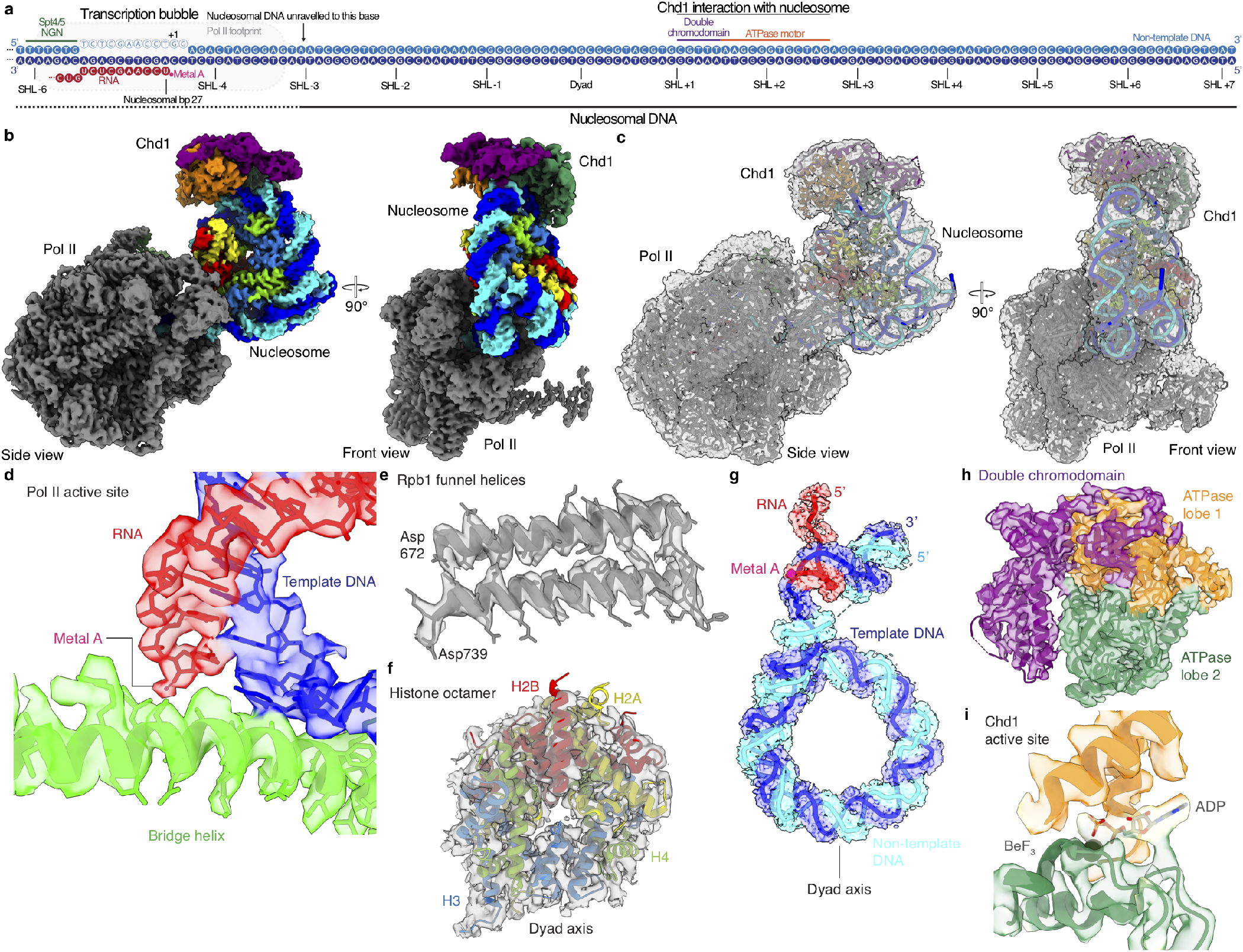
Cryo-EM densities of Pol II-Spt4/5-nucleosome-Chd1 complex. **a**, Protein-nucleosomal DNA contacts of Pol II-Spt4/5-nucleosome-Chd1 complex. Nucleotides are depicted as solid spheres (modelled) or empty spheres (not modelled). SHLs are indicated. **b**, Cryo-EM map (map D) of Pol II-Spt4/5-nucleosome-Chd1 complex. **c**, Pol II-Spt4/5-nucleosome-Chd1 structure with corresponding cryo-EM map (map D). Cryo-EM map is shown in grey. **d**, Active site of Pol II-Spt4/5-nucleosome-Chd1 structure with corresponding density (map D). Metal A is shown as a pink sphere. **e**, Rpb1 funnel helices with corresponding density (map D). **f**, Histone octamer with corresponding density (map D). **g**, Nucleic acids in Pol II-Spt4/5-nucleosome-Chd1 structure with corresponding densities (map D) **h**, Chd1 with corresponding density (map D). **i**, Active site of Chd1 with bound ADP·BeF_3_. ADP is shown in stick representation, BeF_3_ as green spheres. Density from map D.

**Extended Data Fig. 5:**
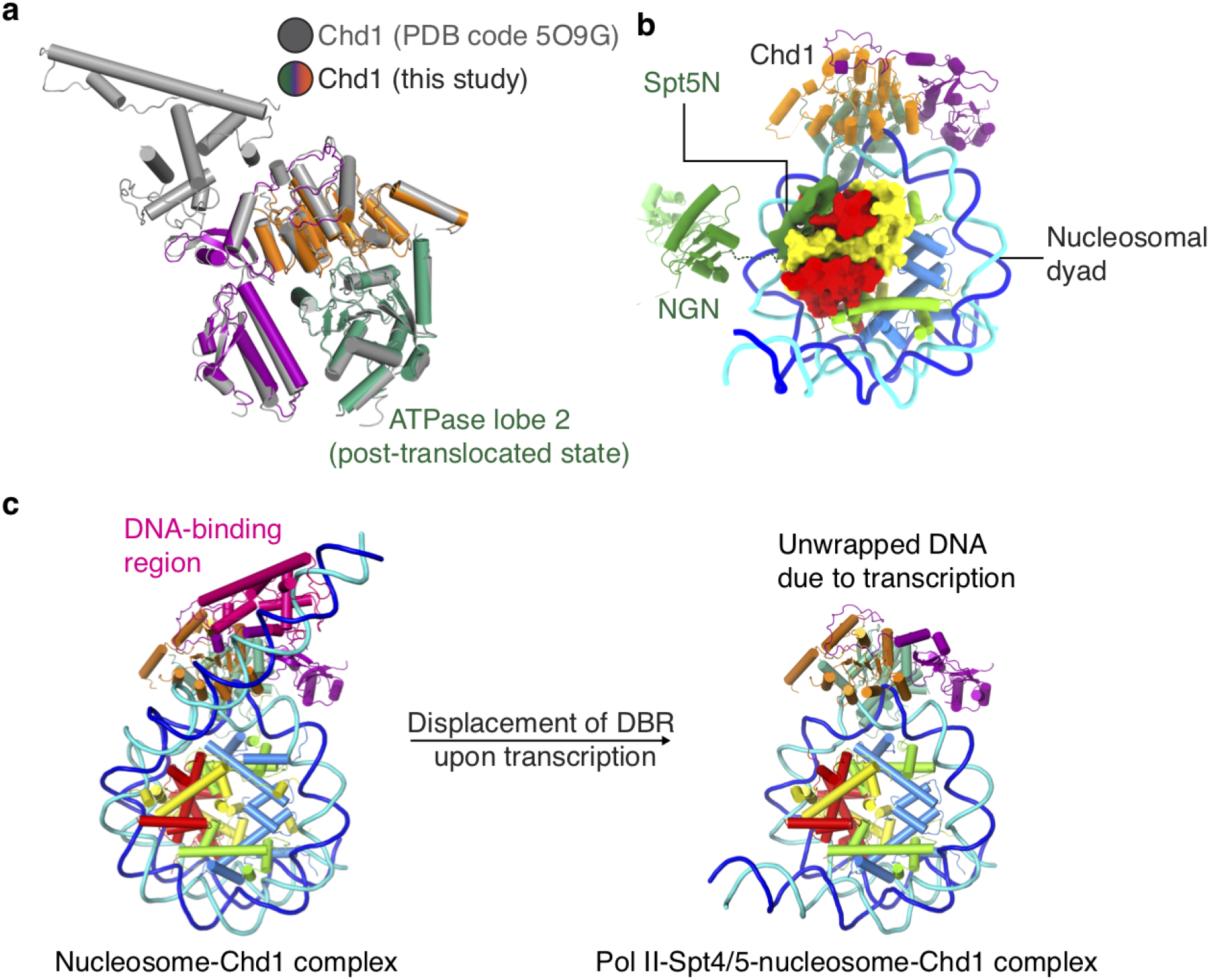
Details of the Pol II-Spt4/5-nucleosome-Chd1 structure. **a**, Comparison of Chd1 (grey, PDB code 5O9G) with Chd1 (this study). The ATPase motor adopts the post-translocated state in both structures. **b**, Density for Spt5 N-terminal region (Spt5N) (low-passed filtered to 9 Å, map D) next to the proximal H2A-H2B dimer (surface representation). **b**, Comparison of a poised nucleosome-Chd1 structure^27^ with the structure of Chd1 bound to the transcribed nucleosome reveals displacement of the Chd1 DNA-binding region (pink) upon transcription.

**Extended Data Fig. 6:**
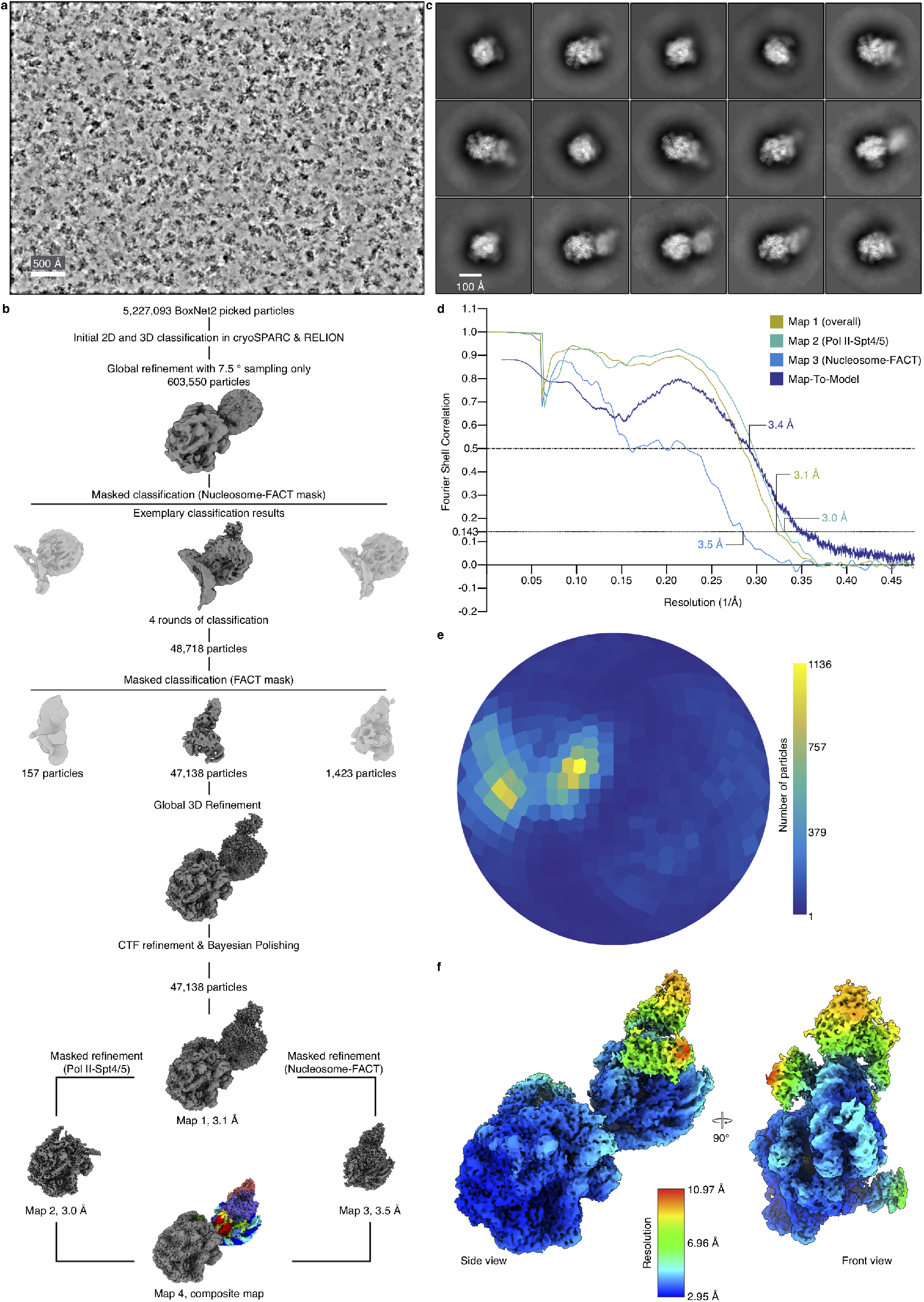
Data acquisition, processing, and data quality metrics for the Pol II-Spt4/5-nucleosome-FACT structure. **a**, Representative denoised micrograph of data collection with scale bar (50 nm). **b**, Sorting and classification tree of Pol II-Spt4/5-nucleosome-FACT dataset. **c**, 2D classes of final refinement show RNA polymerase II and nucleosome-like shape with additional density (FACT) with scale bar of 10 nm. **d**, FSC curves of maps 1-4 and map-to-model. Resolutions at FSC threshold criterions 0.143 and 0.5 are indicated. **e**, Angular distribution of particles employed to reconstruct map 1. **f**, Local resolution of composite map 4.

**Extended Data Fig. 7:**
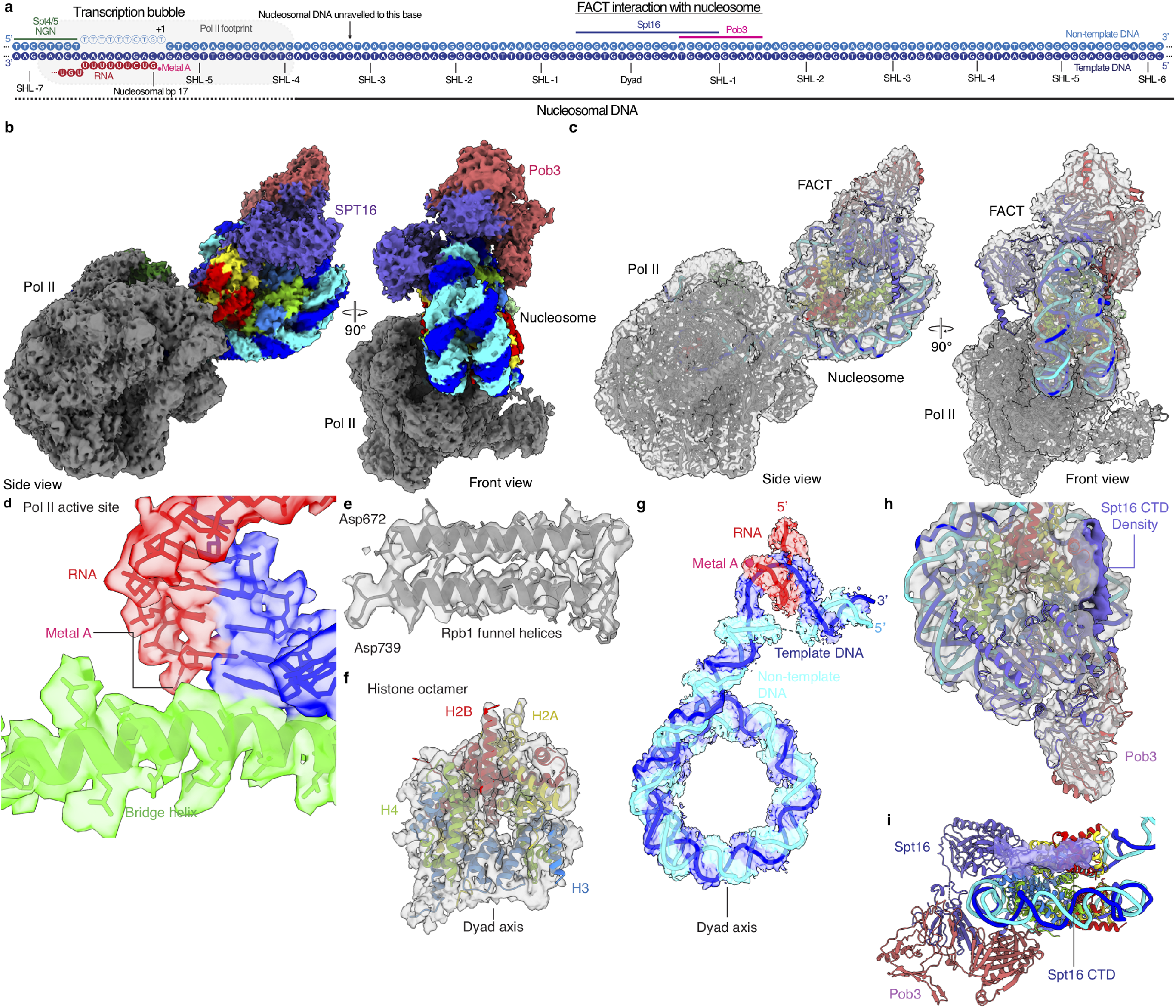
Cryo-EM densities of Pol II-Spt4/5-nucleosome-FACT complex. **a**, Protein-nucleosomal DNA contacts of Pol II-Spt4/5-nucleosome-FACT complex. Nucleotides are depicted as solid spheres (modelled) or empty spheres (not modelled). SHLs are indicated. **b**, Cryo-EM map 4 of Pol II-Spt4/5-nucleosome-FACT complex. **c**, Pol II-Spt4/5-nucleosome-FACT structure with corresponding cryo-EM map (map 4). Cryo-EM map is shown in grey, **d**, Active site of Pol II-Spt4/5-nucleosome-Chd1 structure with corresponding density (map 4). **e**, Rpb1 funnel helices with corresponding density (map 4). **f**, Histone octamer with corresponding density (map 4). **g**, Nucleic acids in Pol II-Spt4/5-nucleosome-FACT structure with corresponding densities (map 4). **h**, Nucleosome with bound FACT and corresponding density (map 4). Density corresponding to the Spt16 CTD is highlighted in purple. **i**, Spt16 CTD density (map4) contacts the proximal H2A/H2B dimer.

**Extended Data Fig. 8:**
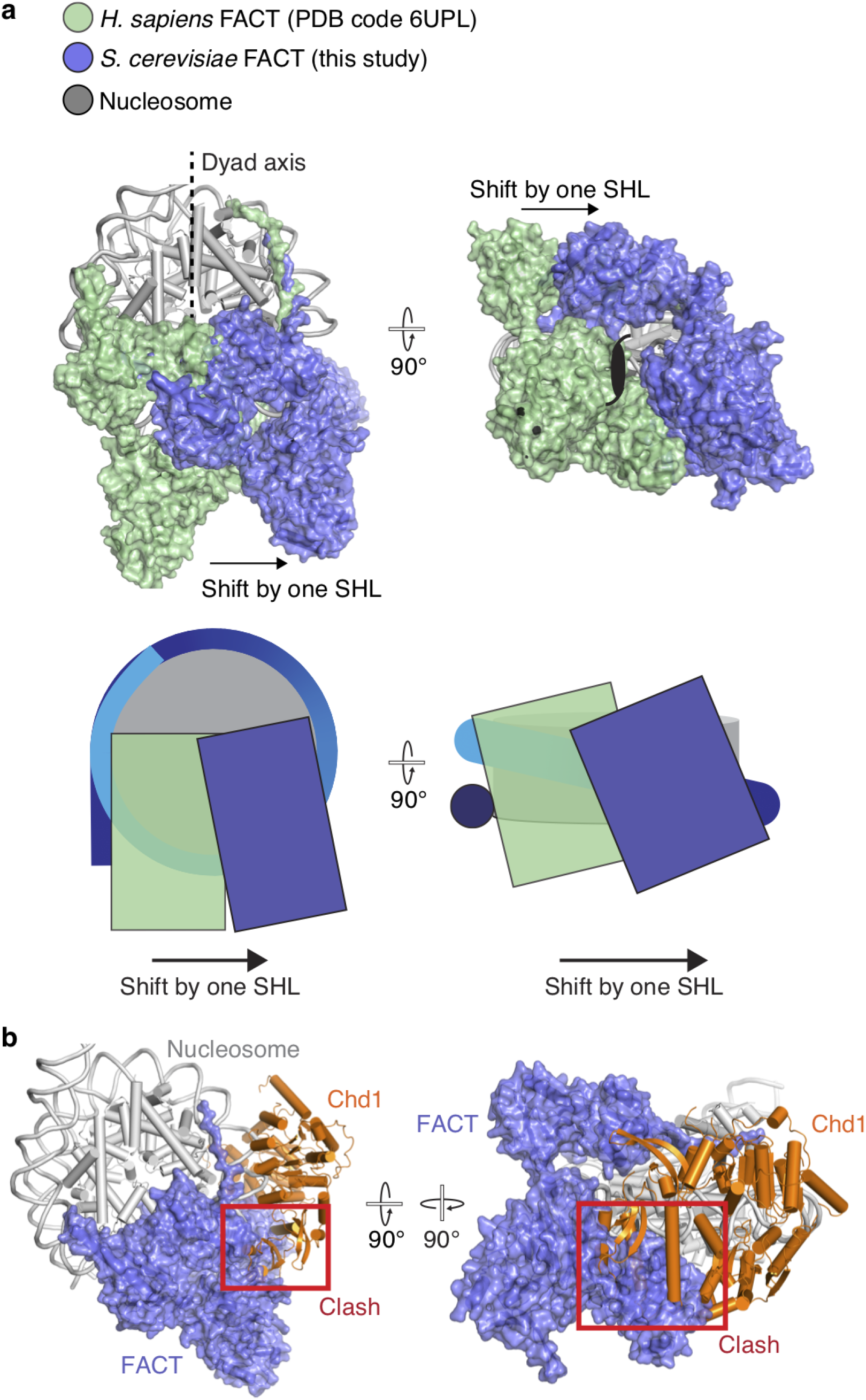
Details of the Pol II-Spt4/5-nucleosome-FACT structure. **a**, Superposition of a subnucleosome-FACT complex (PDB code 6UPL; FACT, pale green) on the Pol II-Spt4/5-nucleosome FACT structures reveals sliding of FACT by one superhelical location. FACT-transcribed nucleosome and the subnucleosome structure were aligned using the histone octamer. Transcribed nucleosome is shown in grey. **b**, Superposition of Chd1 structure with the FACT structure reveals a steric clash between the ATPase lobe 2 and double chromodomain of Chd1 and the Pob3 subunit of FACT.

**Extended Data Fig. 9:**
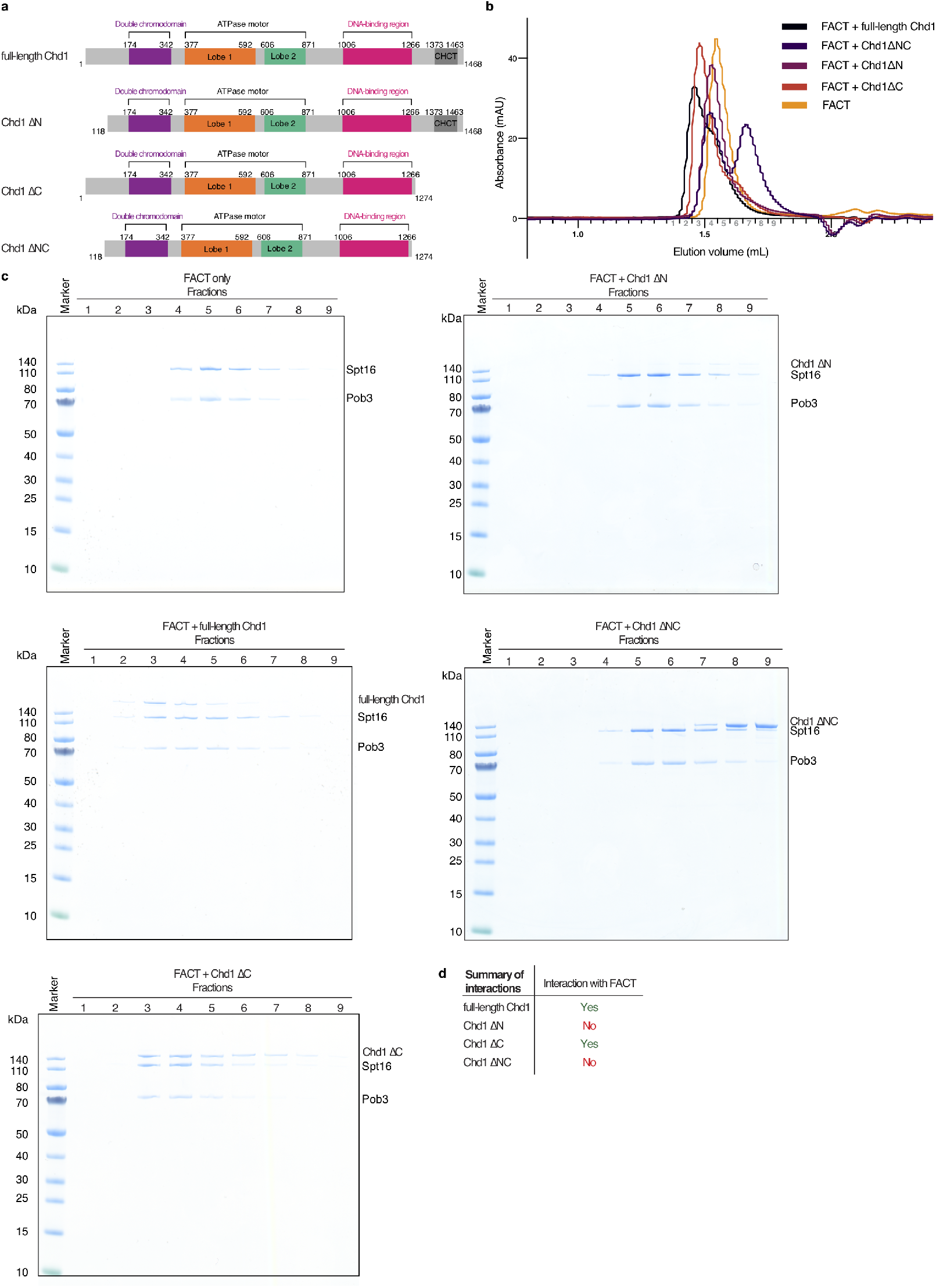
Interaction between Chd1 and FACT. **a**, Domain architecture with different Chd1 constructs; full-length Chd1, Chd1 ΔN (residues 118-1468), Chd1 ΔC (residues 1-1274) and Chd1 ΔNC (residues 118-1274). **b**, Chromatogram of size exclusion chromatography runs to determine regions of Chd1 that interact with FACT. Fractions analysed with SDS-PAGE (c) are indicated with grey numbering. **c**, SDS-PAGE analysis of size exclusion chromatography runs (b) reveals interaction of Chd1 with FACT via the N-terminus of Chd1. **d**, Summary of interaction results.

**Extended Data Fig. 10:**
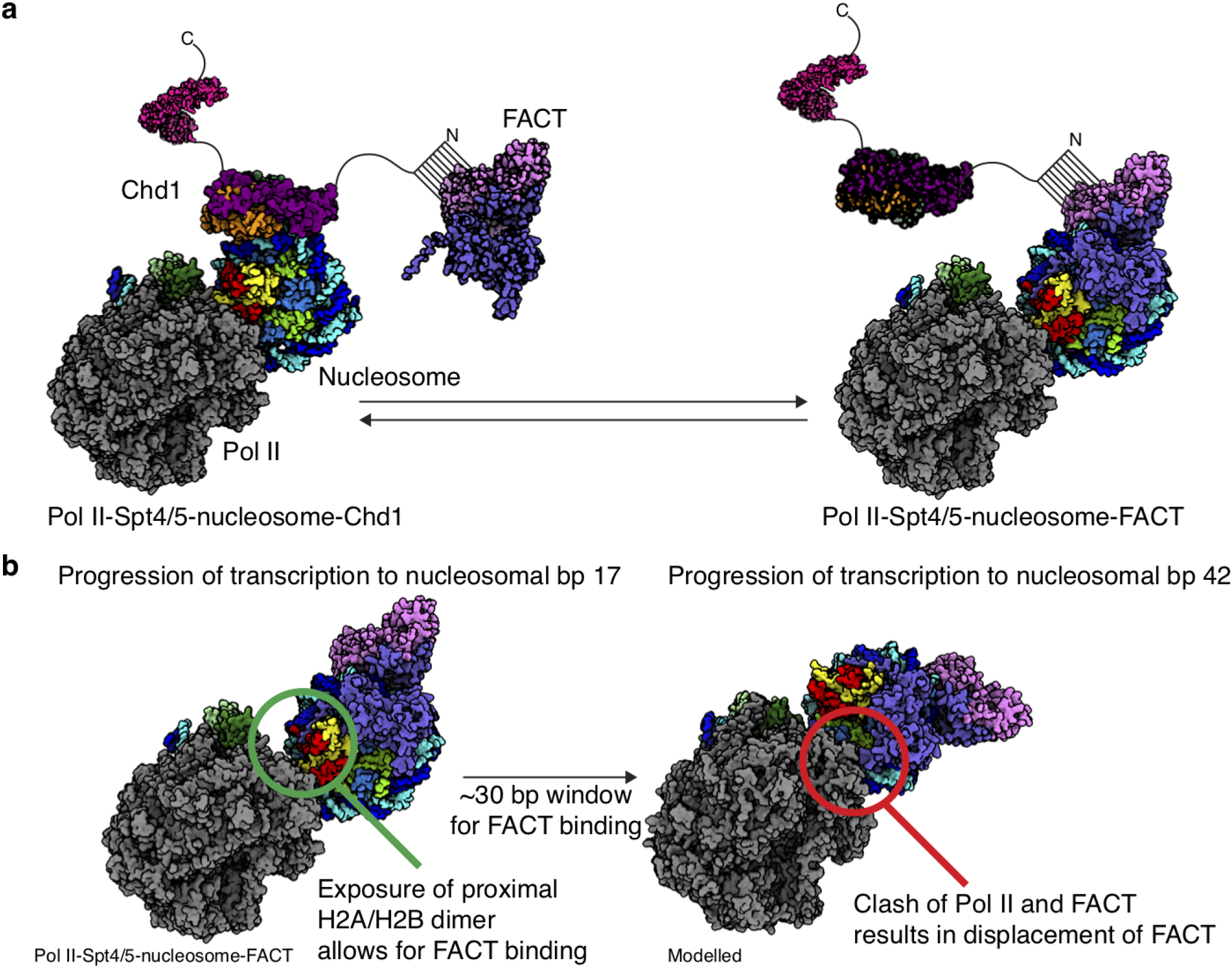
Model for Pol II passage through a nucleosome. **a**, Model for Pol II progression through the proximal part of a nucleosomal substrate. **b**, Structural modelling reveals a 30 bp window for FACT binding during transcription through the proximal part of the nucleosomal substrate.

